# The Pangenome of *Escherichia coli*

**DOI:** 10.1101/2024.06.07.598014

**Authors:** Siddharth M. Chauhan, Omid Ardalani, Jason C. Hyun, Jonathan M. Monk, Patrick V. Phaneuf, Bernhard O. Palsson

## Abstract

Thousands of complete genome sequences for strains of a species that are now available enable the advancement of pangenome analytics to a new level of sophistication. We collected 2,377 publicly-available complete genomes of *Escherichia coli* for detailed pangenome analysis. The core genome and accessory genomes consisted of 2,398 and 5,182 genes, respectively. We developed a machine learning approach to define the accessory genes characterizing the major phylogroups of *E. coli* plus Shigella: A, B1, B2, C, D, E, F, G, and Shigella. The analysis resulted in a detailed structure of the genetic basis of the phylogroups’ differential traits. This pangenome structure was largely consistent with a housekeeping-gene-based MLST distribution, sequence-based Mash distance, and the Clermont quadruplex classification. The rare genome consisted of 163,619 genes, about 79% of which represented variations of 315 underlying transposon elements. This analysis generated a mathematical definition of the genetic basis for a species.

## Background

The first complete genome sequence appeared in 1995 (Fleischmann et al. 1995). Shortly thereafter, the genome sequence of the model *Escherichia coli* K-12 MG1655 strain appeared (Blattner et al. 1997). The genome sequence of a second *E. coli* strain, the enterohaemorrhagic O157:H7 strain, appeared in 2001 (Perna et al. 2001). It had about a 1 Mbp longer genomic sequence than MG1655, encoding about 1000 additional genes representing different traits than those found in MG1655. Following the massive drop in DNA sequencing costs in the late 2000s (Kris A. Wetterstrand 2019 Mar 13), a large number of *E. coli* strain sequences became available (O’Leary et al. 2016; Olson et al. 2023). This data forms the basis for pangenome analysis of the *E. coli* species. In 2013, a study appeared that analyzed 55 *E. coli* genome sequences (Monk et al. 2013). Using metabolic reconstructions and computational systems biology, auxotrophies and colonization sites could be predicted from these genome sequences.

As the number of available genome sequences grew, subsequent studies showed that differential traits between phylogroups could be delineated from sequence and specific pathogenic properties could be deciphered (Fang et al. 2018).

With the availability of low-cost genomic sequencing, strain taxonomic classifications thus moved from phenotypes to genotypes. This started with the creation of the original Achtman multilocus sequence typing (MLST) schema (Wirth et al. 2006). Following this development, the Clermont triplex (Clermont Olivier et al. 2000) and subsequently the quadruplex (Clermont et al. 2013) appeared that deployed PCR assays for discriminating alleles to perform sequence-based phylogrouping. More recently, the whole genome-based Mash distance has been utilized to successfully phylogroup *E. coli* strains, moving the definition of phylogroups to the genome-scale (Abram et al. 2021). Today, the number of *E. coli* sequences in the public domain has reached the 10^5^ scale (Achtman et al. 2022). These sequences contain the full gene complement of these strains. This data availability demands development of novel big data analytic methods that characterize the strains’ genomes based on their full genome-wide gene content.

We can now call the presence/absence of genes across thousands of genomes. The results enable us to form the pangenome matrix (Fang et al. 2018; Norsigian et al. 2018; Seif et al. 2018) for the *E. coli* species. Once formed, this matrix allows us to develop machine learning methods to classify the entire gene complement of these strain sequences into phylogroups. Meaningful classification of strains would allow us to precisely define the genetic basis for differential traits observed between the phylogroups. If phylogroup- and strain-specific traits can be derived straight from sequence, it would reduce the need for strain cultivation in clinical settings and allow for accelerated diagnosis. The full phylogroup definition of the *E. coli* species thus has fundamental and applied implications.

## Results

### Forming the pangenome matrix

We downloaded all available *E. coli* genomes from two public databases, BV-BRC and NCBI RefSeq. This sequencing data was subjected to quality controls and admissions criteria from pangenomic studies (**Fig. 1a**, Methods). The result was a collection of over 10,000 high-quality genome sequences, of which 2,377 were high-quality complete sequences that were used for pangenome analysis. These sequences were collected from a wide variety of isolation sources, including humans, land animals, and various species of birds (**Fig 1b**). Most of the strains did not contain any plasmids, with notable exceptions (e.g., a Phylogroup G strain containing seven plasmids) (**Fig 1c**). We call this curated collection of sequences and its resulting pangenome a **G**enome **E**ncyclopedia of **N**otable **O**bserved **Mi**croorganisms **C**urated for **U**niversal **S**tudy (GENOMiCUS).

**Figure 1:**
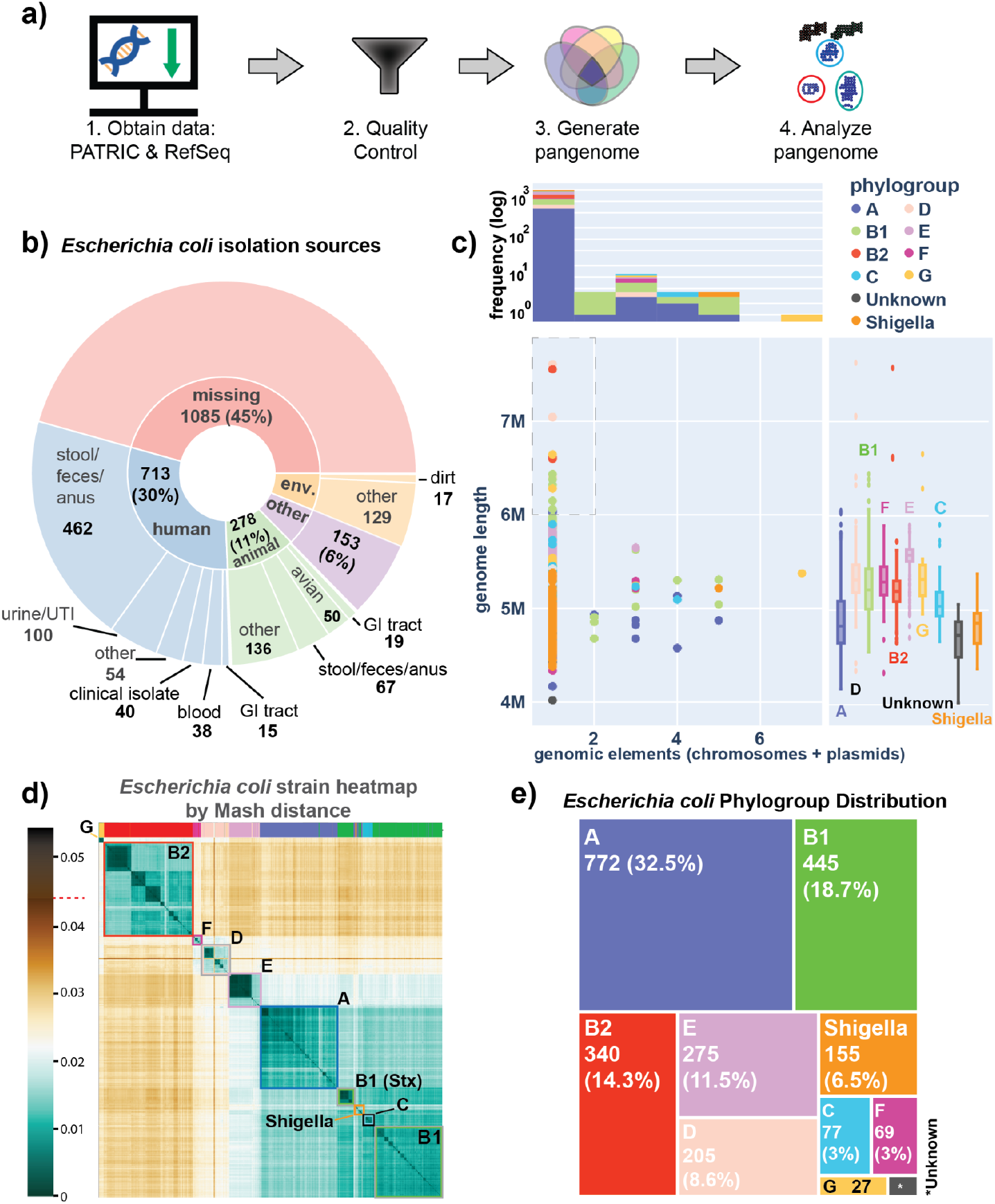
Processing and classification of a 2,377 complete Escherichia coli genome compendium (GENOMiCUS) **a)** The workflow used in this study. Genomes were downloaded from PATRIC (now BV-BRC) and RefSeq, after which they were deduplicated and filtered based on their quality metrics (see **Methods**). The resulting 2377 complete genomes form a high-quality compendium of strains for detailed pangenome analysis. We call this compendium the Genome Encyclopedia of Notable Observed MIcro-organisms Curated for Universal Study (GENOMiCUS). **b)** A sunburst plot showcasing the different isolation sources for the bacteria in this compendium. While most of the 1332 isolation-site-annotated strains come from humans (713), there are many strains isolated from animals (278) and various other environmental niches (146). **c)** Scatterplot summarizing properties of the genomes by genome length (y-axis) vs. number of genomic elements (chromosomes + plasmids) (x-axis), colored by phylogroup as calculated in-silico by the ClermonTyping github package (Beghain et al. 2018). Note that many Shigella strains were incorrectly classified by ClermonTyping as belonging to Phylogroup A, and so any strains which were known to be Shigella were manually separated into a separate class for better identification. 19 strains were found to have a genome size greater than 6 Mb. 16 of those 19 strains were clinical isolates from ICDDR,B from patients who had diarrhoeal disorders. Above the scatterplot is a histogram showcasing the genomic element distribution within the strains of the pangenome, also colored by phylogroup. Note: in this context, a “genomic element” refers to both the main chromosome and any additional plasmids found in the organism. To the right of the scatterplot are phylogroup-specific boxplots describing the distribution of genome lengths per phylogroup. **d)** A heatmap of the pairwise Mash distances for all 2,377 E. coli strains of GENOMiCUS based on sequence analysis. Distances range from 0 to 0.04 and the highest Mash value (0.044) is denoted with a red dash on the colorbar Note that a pairwise Mash distance of 0.05 equates to an average-nucleotide-identity (ANI) of 95%, both of which correspond to a 70% DNA-DNA reassociation value, the historical definition of a bacterial species (Konstantinidis and Tiedje 2005; Ondov et al. 2016). The highlighted bars at the top of the heatmap identify the Mash-based clusters of this compendium. Phylogroups are annotated on the heatmap, showing the correspondence between these phylogroups and the Mash-based clusters. **e)** Treemap illustrating the distribution of E. coli strains by phylogroup as calculated in-silico by the ClermonTyping github package (Beghain et al. 2018).

Genomes can be classified using sequence characteristics. The Mash distance between genome sequences has been shown to quantify their differences (**Fig 1d**) (Ondov et al. 2016; Abram et al. 2021). One can now cluster a series of genome sequences based on global sequence similarity. A heatmap classification of the sequences used in this study shows that Mash distances lead to phylogroup classification, consistent with a previous study (Abram et al. 2021). In addition, phylogroup designation based on the Clermont quadruplex standard can be computed from the sequences (summarized **Fig 1e**) (Clermont et al. 2013; Beghain et al. 2018). It shows that phylogroups A, B1, and B2 had the highest number of strains in the collection analyzed. In contrast, the recently defined phylogroup G (Lu et al. 2016; Clermont et al. 2019) had relatively few complete strain sequences available for analysis.

### Stratifying the pangenome into three categories of genes

A pangenome can be stratified into three main categories of genes:

- The **core genome** consists of the genes found in all, or nearly all, of the strains. These genes, therefore, can be taken to define the species. For the collection of strain sequences analyzed here, the core genome consists of 2,398 genes, 80% of which have known functions.
- The **accessory genome** is composed of 5,182 genes. These are genes that are found in many, but not all strains. The accessory genes, being variably present, can be used to define the gene portfolio of the phylogroups, as described below.
- The **rare genome** consists of genes unique to a strain or found in a relatively small number of strains.

These three categories of genes are deciphered from the frequency of gene occurrence in the collection of strain sequences (**Fig 2a**). This gene frequency histogram shows the number of genomes containing a particular gene. Taking the cumulative sum of the gene frequency, we get a cumulative gene distribution that is used to formally determine the boundaries for the core, accessory, and rare genomes (**Fig 2b**) (Hyun et al. 2022).

**Figure 2:**
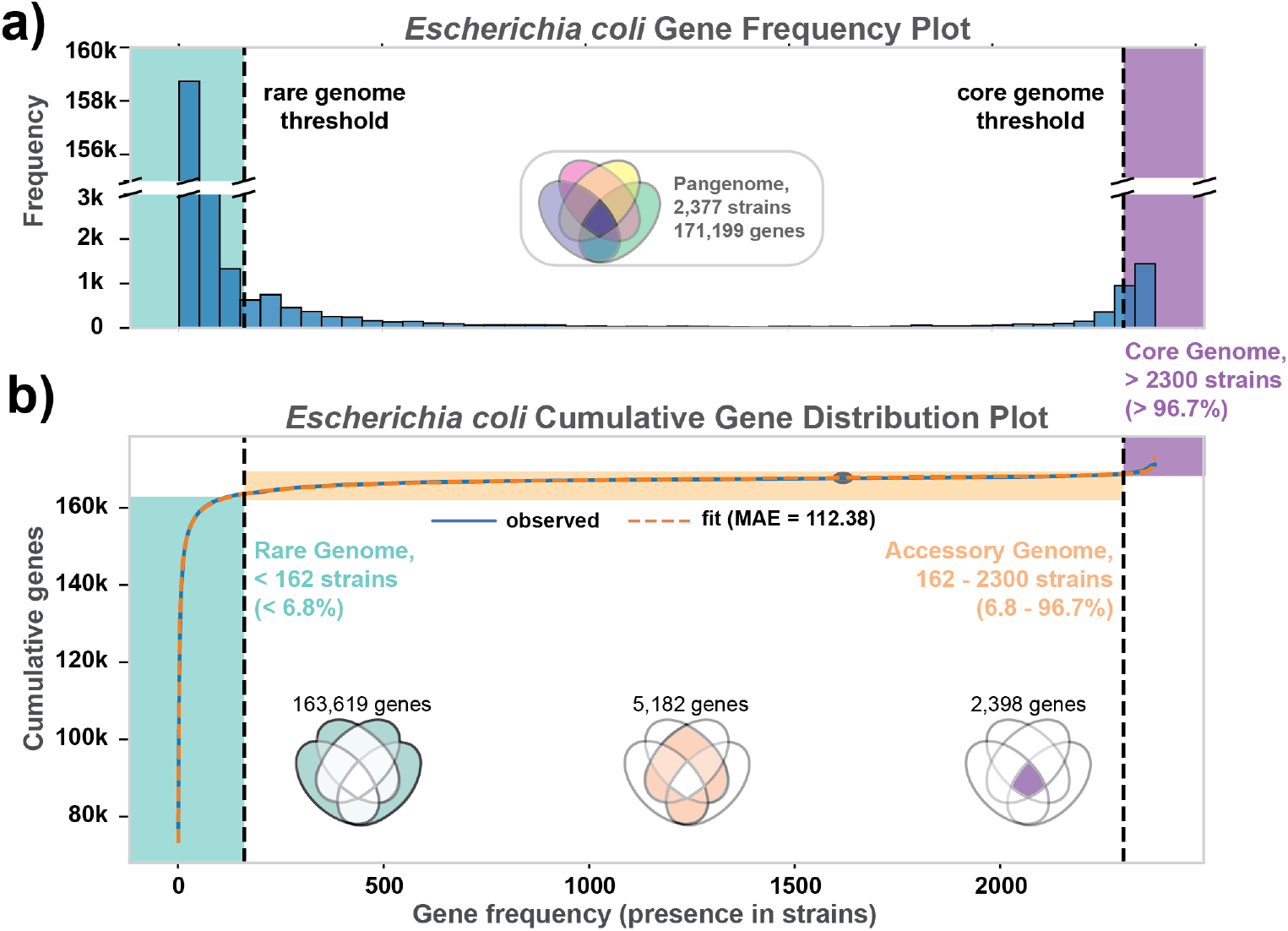
Global distributions of gene frequencies and functions in the Escherichia coli pangenome. **a)** Gene frequency distribution across the 2,377 curated genomes in GENOMiCUS. Genes present in all 2,377 strains appear at the histogram’s right end. Progressing leftward, subsequent bars show genes found in nearly all strains, decreasing in frequency, until reaching genes unique to just one strain at the extreme left. **b)** The cumulative gene distribution function (Hyun et al. 2022). The gene frequency distribution was fitted to a double-exponential form (with median absolute error or MAE = 176.31) and the inflection points determined. Based on these inflection points, the genes in the pangenome were divided into the core (comprising 2,398 genes), accessory (comprising 5,182 genes), and rare (comprising 163,619 genes) genomes (See Methods).

### Defining the genes in the core, accessory, and rare genomes

The boundary between the core and accessory genome separates (near-)omnipresent genes from variably present accessory genes. Defining the boundary between the rare and the accessory genome is more subjective. The definition of these boundaries in this study is described in **Methods**, and they lead to the identification of 2,398 core genes, 5,182 accessory genes, and 163,619 rare genes. The exact definition of these boundaries does not affect the major conclusion of this study (see **SI**).

The number of genes classified into the core genome can be plotted with the number of strains considered (Heaps’ Plot, **Fig S1**). This curve levels off fairly quickly with the number of genomes considered, and stays flat at 2,398 genes, defining a closed core genome. The number of genes classified as accessory genes similarly levels off at 5,182 genes. This observation shows that the accessory genome is also closed. The closed nature of the accessory genome makes it possible to analyze the phylogroup gene content in novel and mathematically rigorous ways, as shown in the next section.

Thus, after a certain number of strains, the discovery of novel genes in the pangenome is driven by occurrence of rare genes; the median number of such genes per strain is 270, and 163,619 total amongst the 2,377 genomes studied. These rare genes will confer unique characteristics onto the strain in which they reside.

### Traits in the core genome

The genes common to the 2,377 strains represent the core genome. Thus, there is a uniform genetic basis for certain traits. For instance, the core genome contains 18 of 29 two-component systems, consistent with previous findings (Rajput et al. 2021). One of 68 biosynthetic gene clusters, two of 130 AMR genes (*lnt*, an apolipoprotein N-acyltransferase and *narP*, a nitrate/nitrite response regulator), 382 of 127,223 transposable elements, and 21 of 7,925 motility genes (pili, fimbriae, flagella, and supporting proteins) are in the core genome. There are 1,006 metabolic genes in the core genome, which is slightly higher than the 976 genes previously reported (Monk et al. 2017) (**Fig S2**). There are still 462 genes of unknown function (y-genes) in the core genome, comprising 19% of all core genes.

### The accessory genome has a clear mathematical structure

The accessory genome is effectively closed (**Fig S1**), enabling a comprehensive analysis of gene-level diversity amongst the 2,377 strains. To do so, we form the **P** matrix (genes x strains) for just the accessory genome. This **P** matrix can be decomposed using non-negative matrix factorization (NMF) (Lee and Seung 1999; Devarajan 2008) to define the genes that belong to strains of a particular phylogroup. NMF factors **P** into two matrices:

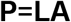

**L**, whose columns consist of gene weightings that define a *phylon* (the genes common amongst similar strains, often belonging to the same Clermont phylogroup and/or MLST cluster), and **A**, whose rows give a strain’s *affinity* (or closeness) of a genome to a phylon (**Fig 3a**). The column space of **L** is a convex cone, as all its values must be non-negative. Each column of **L** (a phylon vector) represents an edge of a polygon (**Fig 3b**).

**Figure 3:**
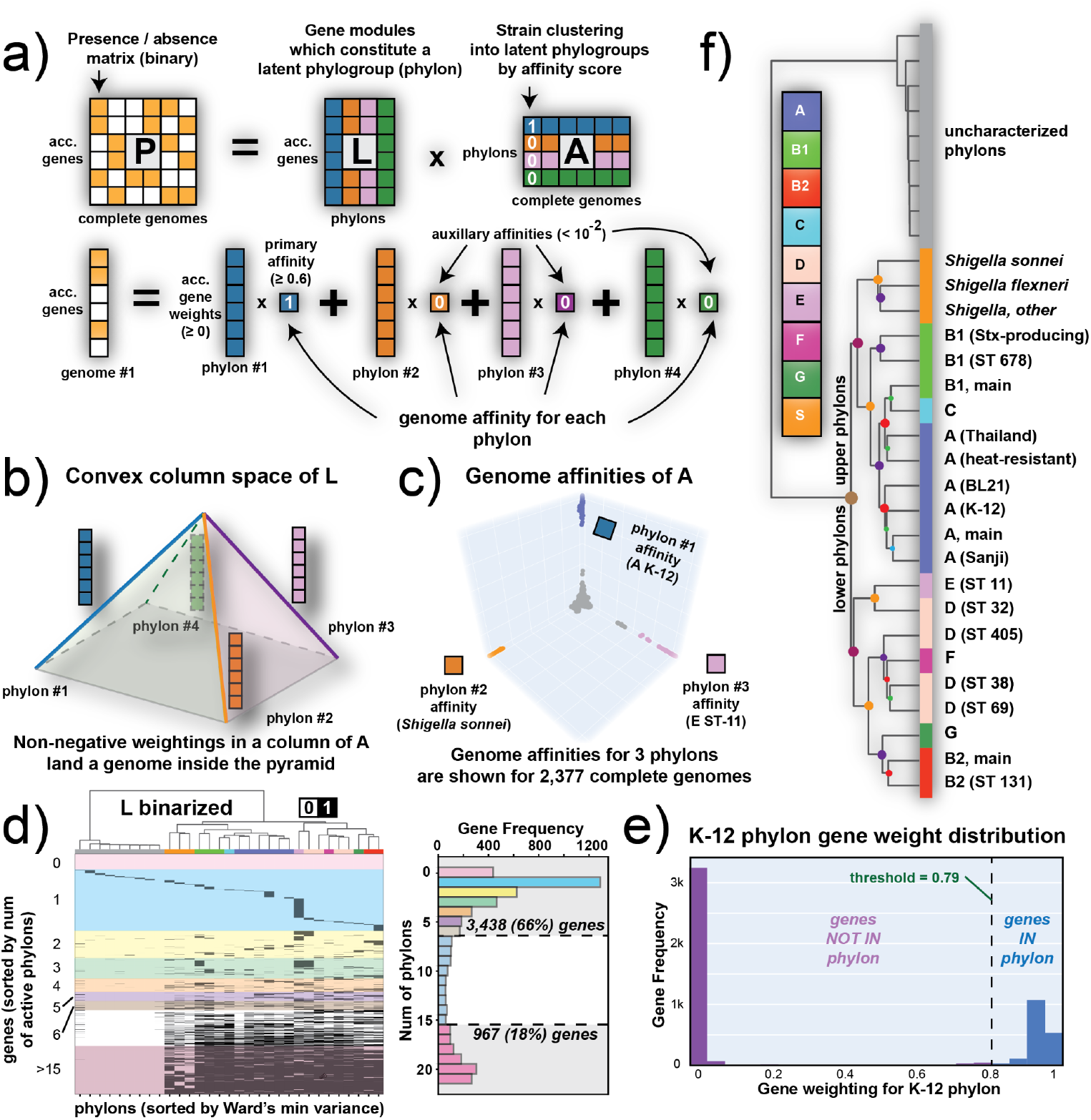
The fundamental mathematical structure of the E. coli accessory genome. Characteristics of the NMF decomposition of the pangenome matrix P. **a)** A column of **P** (i.e., genome #1) is a linear combination of the phylon vectors as determined by the weights in the corresponding column of **A. b)** Since the phylon vectors are non-negative, they span a polygon as its edge vectors. A positive linear combination of the **l**_i_ vectors lands inside the polygon. **c)** Since there is typically only one dominant value in a column of **A**, the reconstruction of a column in **P** (i.e., one genome) lies close to a phylon vector (i.e., the edges of the polygon) as is evident for the 2,377 sequenced strains used. **d)** A clustermap of the binarized **L** matrix. Colors on top correspond with classically-defined phylogroups as determined by ClermonTyping. Columns are clustered using Ward’s minimum variance method, and rows are sorted by gene frequency in each phylon (i.e., genes in zero phylons are at the top, genes in 22 phylons are at the bottom). The dendrogram at the top of **L**, showing the clustering of its columns, is the same as that used in panel **f**. In this graphical representation the black elements designate that the gene responding to that row is found in the phylon that the column represents. White elements mean that the corresponding gene is not found in the phylon. The histogram to the right of the clustered **L** matrix showcases the gene frequency across multiple phylons (i.e., how many phylons a gene is present in). The colors in L-binarized correspond to the colors on this histogram and showcase the distribution of genes by their number of active phylons. 3,438 (66%) of the 5,182 accessory genes are found in six or fewer phylons, with the plurality being genes active in only one phylon (1,289 single-phylon genes, 25% of all 5,182 accessory genes). **e)** A gene weight distribution for one particular phylon consisting of K-12 strains in the **L** matrix. Most genes have a weighting close to zero, with a notable cluster having weightings between 0.8 and 1. The genes with low weightings (below the threshold indicated by the dashed line) are binarized to zero and considered not to be part of the phylon while genes with high weightings are binarized to one and considered to be constituents of this phylon. The threshold for binarization is determined for each phylon using k-means clustering (see **Methods**). **f)** A dendrogram of all 31 phylons based on clustering the binarized **L** matrix shown in Panel **d**). The uncharacterized phylons are separated, mainly consisting of phage genes and other mobile elements.

NMF gives a clear mathematical description of the gene portfolio of a phylogroup found in the *E. coli* pangenome. The gene list found in all strains of a phylogroup is given by a phylon, or a column in **L**. Few strains will correspond perfectly to a phylon as its gene list may differ slightly from that given by the columns in **L**. The affinity matrix, **A**, shows how close a strain is to a phylon as the elements in a column in **A** give the phylon composition of a particular strain. This feature is demonstrated with the color coding of the matrices in **Fig 3a**. The 3D image of the location of all strains relative to three of the columns of **L** is shown in **Fig 3c** for all 2,377 genomes in this study. Strains of a phylogroup are close to one of the phylon vectors shown (i.e., high affinity for the phylon), while the rest of the strains that are not in these three phylogroups are close to the origin (i.e., low affinity for these phylons).

Almost all strains have a dominant phylon as they lie close to one edge of the column space. Thus, the affinity scores in the column of the **A** matrix that corresponds to a particular strain places each genome inside the convex solution space. Most of these affinities are small and close to zero, typically with only one dominant affinity per genome, revealing that most strains reside close to the edges of the convex space. An image of the binarized form of **A** is shown in (**Fig S4**).

### Biological meaning of the pangenome’s mathematical structure

The columns of **L** show that NMF breaks **P** into the eight classically-defined phylogroups (plus Shigella, see **Methods**) and sub-phylogroups thereof (**Fig 3d**). There are 22 of these columns, and then an additional nine unclassified columns of **L** that represent mobile elements (see below). The **L** matrix shown in **Fig 3d** is binarized. The weightings are close to unity (gene in phylon) or zero (gene not in phylon), as shown in **Fig 3e**.

Thus, the NMF decomposition of **P** for the accessory genome reveals phylons defined by their list of genes. It also shows how each strain’s gene set maps onto these phylons. NMF segregates genes to a phylon concordant with previous phylo-grouping methods of strains in *E. coli* (**Fig 3f**): the Mash distances (**Fig 1d**), the Clermont quadruplex, and the MLST typing (see **Table S5**). NMF allows us to go from differential traits between phylogroups to their genetic basis.

### The *E. coli* pangenome consists of two distinct groups of phylons

The phylons are divided into two major groups (**Fig 3f**). One group, which we collectively call the lower phylons – G, B2, D, F, and E – are genetically dissimilar to the strains found in the upper phylons that correspond to phylogroups A, B1, C, and Shigella. This is the first split in the hierarchical phylon clustering tree. Of the 5,182 genes found in the accessory genome, 765 are found exclusively in strains of the upper phylons, with 1,244 found exclusively in strains of the lower phylons (**Fig 4**). While some of these genes have no known function/ortholog, many do. In fact, 98 of the 765 upper-phylon-exclusive genes have a known metabolic function, with an additional 34 having a known motility function. Similarly, 213 of the 1,244 lower-phylon-exclusive genes are metabolic in nature, with 73 having motility functions. The distribution of genes in many other functional categories (such as transcription factors, metabolic functions, pili, motility, membrane-, and phage-genes) are highlighted in **Table S2**. The columns of **L** thus gives us detailed information about the differential gene contents of the phylons, and thus gives the basis for finding the genetic basis for differential traits between phylogroups.

**Figure 4:**
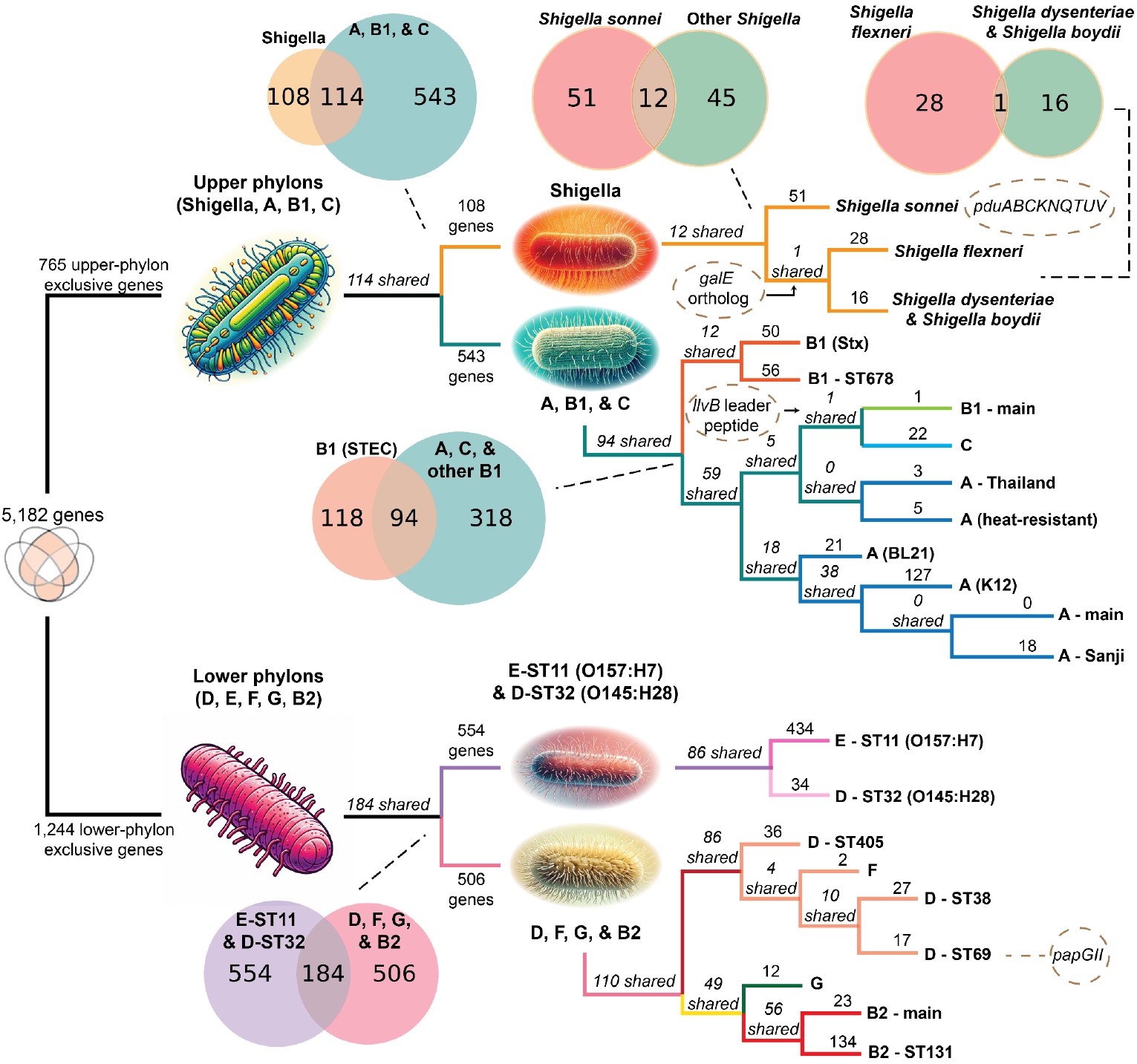
A clustering diagram of the phylons (see Fig 3f) that highlights the groups of exclusive genes that follow one branch and not the other at each branch point. The numbers above the line leading to a split indicate exclusive genes (i.e., genes found in one group of phylons but absent in the other). Numbers in italics specifically indicate shared genes that are found across all groups. The Venn diagrams showcase the same information for particular branch points. The function and identity of special genes of interest are discussed in the main text, and detailed in **Table S2**. Four specific genetic traits of interest are highlighted in dashed ovals, such as the papGII operon to phylon D (ST69). This sequence variant of papG in this operon is associated with UTIs that can become bacteremic (Biggel et al. 2020; Cuénod et al. 2023)

A further examination of the metabolic gene content between these two distinct groups of phylons reveals that the upper-phylon-exclusive genes code for glycosyltransferases, exo-alpha-sialidases (glycoconjugates), beta-glucosidases, beta-glucuronidases, hyaluronidase, and monogalactosyldiacylglycerol lipases (monogalactosyl-diacylglycerols), among others. These enzymes are all involved in processing and digesting glycans in a host, including sialic acids. Sialic acids and their derivatives often form the ends of glycans on various glycoproteins and glycolipids which coat the surfaces of most vertebrate and bacterial cells. They are known to behave as a signal to specific bacteria upon reaching a vertebrate environment suitable for colonization (Varki and Gagneux 2012; Schauer and Kamerling 2018). The presence of these enzymes in the upper phylons suggests that Shigella, Shiga-toxin producing *E. coli* (STEC), and other pathogenic strains in phylogroups A, B1, and C have the ability to identify and process these sugar moieties, which may enable them to better colonize their hosts.

A parallel inspection into the lower-phylon-exclusive metabolic genes reveals they code for various bacterial capsule formation proteins, Amadori product degradation (specifically fructoselysine / psicoselysine degradation), as well as three distinct fructose-bisphosphate aldolases, one of which is GatY (Fang et al. 2018). All of these are known proteins found in tagatose-competent *E. coli* strains (primarily found in phylogroups B2 and D) which can utilize these enzymes to digest the glycans in the mucus within the human GI tract(Fang et al. 2018).

The secondary splits in the phylon clustering tree allow further tracing of genes and thus segregation of the genetic basis for traits. Three further splits are discussed here. A full description of the classification tree requires a comprehensive study that will result in the full genetic definition of *E. coli* and all its phylogroups.

#### Shigella strains exhibit gene gains as well as gene losses

Continuing the segregation of genes down the tree of phylons, as defined by NMF (**Fig 3f, Fig 4**), we see that the upper phylons split by gene content into Shigella strains and those that belong to the classically defined A, B1, and C phylogroups. A closer look at this split reveals that the strains in the Shigella phylons contain 108 exclusive genes not found in the A, B1, and C strains, in addition to not containing 543 genes found in these three classically-defined phylogroups. This suggests that Shigella has undergone both gene gain and loss during its restriction to human hosts and adaptation to the human intestinal mucosa (The et al. 2016). For example, in the *nadA* and/or *nadB* genes encoding the enzyme complex that converts L-aspartate to quinolinate, a precursor to NAD resulting in nicotinic acid auxotrophy is lost in these strains (Di Martino et al. 2013). However, other genes are gained, including nine genes that form the propanediol utilization (*pdu*) operon. Propanediol is produced when fucose (a component of mucin) is metabolized under anaerobic conditions (Dogan et al. 2014). Of particular note is the *pduC* gene in this operon, which is enriched in adherent-invasive *E. coli* found in the microbiome of Crohn’s disease patients (Viladomiu et al. 2021). Note that *Shigella sonnei* is separated from the rest of the Shigella strains, which all contain a *galE* ortholog that *Shigella sonnei* itself lacks (**Fig 4**).

#### Pathogenic A strains are more closely related to B1 and C strains than to commensal A strains

The upper phylons split into Shigella strains and strains in A, B1, and C phylogroups. The A, B1, and C strains further split with Shiga-toxin producing *E. coli* strains (B1-Stx and B1-ST678) forming their own subgroup. Interestingly, the next split between these strains in the other branch occurs between commensal A strains and B1, C, and pathogenic A strains.

Specifically, these pathogenic A strains are those that came from foodborne illnesses in Thailand (A-Thailand) and those found to be heat-resistant in meat (A-heat-resistant). The commensal A strains are heavily used in laboratory work and in biomanufacturing (specifically the K-12 MG1655 strain).

#### Lower phylons have a subgroup containing *E. coli* O157:H7 and O145:H28 strains

The lower phylons split into two subgroups, with E-ST11 (O157:H7) and D-ST32 (O145:28) strains separating from the other group of lower phylons (see **Fig 4**). These two serovars (O157:H7 and O145:H28) are known to have shared a common evolutionary lineage (Cooper et al. 2014). This shared lineage is directly reflected in their shared gene content, with 86 shared (accessory) genes between them. Of these genes, 33 have no known function, while 11 of them are metabolic, 10 are motility-related, and 9 are transcription factors. The 11 metabolic genes primarily code for various transporters, oxidoreductases involved in glycolytic pathways, and a class-II fructose-bisphosphate aldolase. Furthermore, all motility genes code for fimbrial gene orthologs of the yadCKLM-htrE-yadVN operon. This operon is cryptic under normal laboratory conditions but when constitutive expression is induced, it promotes biofilm formation in minimal media on a variety of abiotic surfaces and produces surface fimbrial structures (Korea et al. 2010). Constitutive expression of this operon also results in increased adhesion of cells to xylose-rich glycans, increased adherence to intestinal epithelial cells, and can also modulate the inflammatory response of host cells (Larsonneur et al. 2016).

#### D and F strains are very closely linked, as are G and B2 strains

The remaining lower phylons consist of D, E, F, and B2 strains. These strains cluster into two distinct groups: the first group consists of D and F strains while the second group consists of G and B2 strains. The first cluster shares 86 genes, and D-ST405 strains appear to be the most distinct, even more so than F strains. This clustering suggests that the classically-defined D and F phylogroups of *E. coli* are more closely related to each other genetically than previously thought. This is similarly the case for G strains, which were already known to be more closely related to F and B2 strains than others (Clermont et al. 2019).

#### Uncharacterized phylons contain mobile genetic elements

The same mobile elements can be found in strains of many phylogroups. Remarkably, NMF detects this characteristic and factors out these mobile genes into a set of nine ‘uncharacterized’ phylons since they can be columns in the addition that forms the gene set in a strain, see **Fig 3b**. These mobile elements are described in **Table S2** and include sex pili, F-plasmid operons, and various phage genes, among others. The mobilome o*f E. coli* is 9-dimensional.

### Traits found in the rare genome

The rare genome consists of 163,619 genes (**Fig 2b**), of which 127,223 (or 79%) are transposable elements (TEs) (**Fig S5a**). These TEs fall into 315 unique categories of TEs. The number of the 40 most frequent TEs and the number of passenger genes (a.k.a cargo genes) they carry is shown in **Fig 5a**. About 3% of the most frequent TEs carry 773 unique passenger genes, that with replicate occurrences, give a total count of 3,631 rare genes. These 3,631 genes fall into 24 functional categories, of which the largest COG category is unknown function (S), followed by energy production and conversion (C) and transcription (K) (**Fig S5b**). Thus, the genetic diversity of the rare genome is effectively much narrower than its raw gene count indicates.

**Figure 5.**
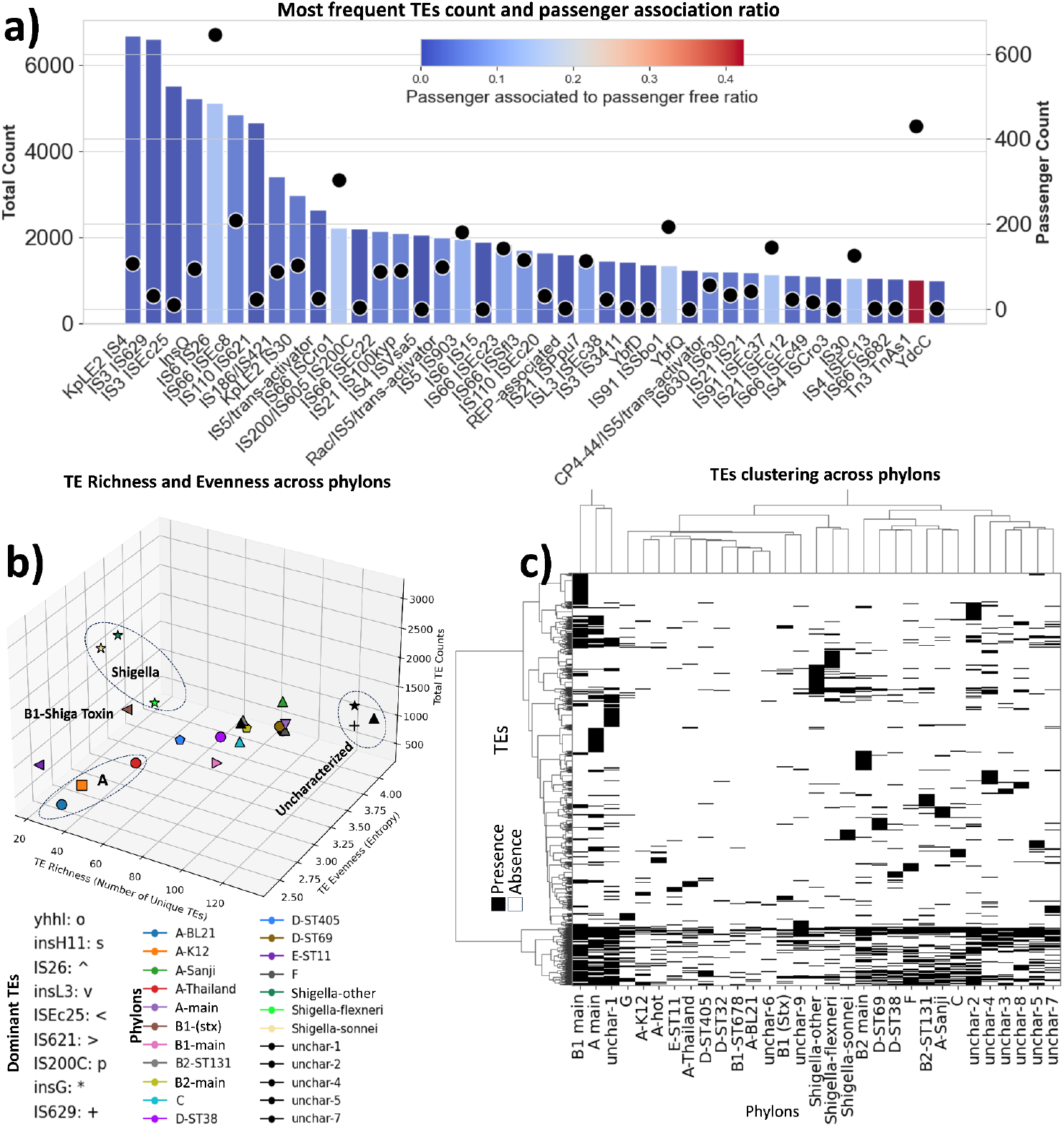
Transposable elements (TEs) in the rare genome. **a)** Frequency of the 40 most abundant TEs of the 315 TE types found in the pangenome, and the ratio of passenger-free to passenger-associated TEs. Bar plot represents count of each group of TEs (top x-axis), bar color indicates the ratio of passenger free-TEs to passenger-associated TEs, and dots indicate the count of passenger genes associated with each group of TEs (bottom x-axis). Naming convention of the TEs is derived from PROKKA annotation. **b)** Phylon TEs count, richness, and entropy; colors represent phylons, signs indicate the dominant TE found in each phylon, TE richness shows how many unique types of TE inhabited a phylon, TE evenness shows how evenly a phylon is infected with different types of TE, higher values of evenness indicates greater entropy. Phylons with fewer than 30 genomes were excluded from the analysis. For phylons with more than 30 genomes, a random subset of 30 genomes was selected, and the remaining genomes were excluded (Fig S5c illustrates the sensitivity of TE richness, evenness, and count metrics to the random sampling of genomes); the symbols on the plot represent the dominant TE in each phylon: circle (‘o’) for yhhI, square (‘s’) for insH11, upward triangle (‘^’) for IS26, downward triangle (‘v’) for insL3, left-pointing triangle (‘<‘) for ISEc25, right-pointing triangle (‘>‘) for IS621, pentagon (‘p’) for IS200C, star (‘*’) for insG, and plus (‘+’) for IS629. **c)** Represents a heatmap generated using CD-HIT clustering results, depicting the distribution of TEs across various phylons. Each cell in the heatmap represents the presence (black) or absence (white) of a specific group of TEs in a given phylon. The hierarchical clustering, employing the Ward method, is applied both horizontally and vertically, illustrating the grouping of similar TEs and phylons, respectively. The dendrograms adjacent to the rows and columns indicate the clustering relationships.

In the Tn3 family, particularly within the *TnAs1* transposase family, a notably higher ratio of passenger-associated TEs (42%) was observed compared to all other TEs categories; interestingly, studies have identified the Tn3 family of transposons as a factor in the recent surge of carbapenem and colistin-resistant Enterobacteriaceae (Cuzon et al. 2011; Nordmann et al. 2012; Borowiak et al. 2017; Zhang et al. 2019). The richness (representing the number of distinct types of TEs found in a phylon) and entropy (indicating the degree of unevenness in the distribution of various transposable element types within a phylon, with higher values suggesting a less uniform population) of TEs are greater in uncharacterized phylons, indicating higher diversity and greater heterogeneity in the distribution of TEs (**Figs 5b and 5c**). In three out of the four phylons related to phylogroup A, comparatively low levels of TE entropy, richness, and count were observed. This result suggests that reduced replicative activity of TEs is exhibited within these phylons. Furthermore, these phylons are characterized by the dominance of a specific TE type, namely *yhnl* for A-Thailand and A-BL21, and *‘insH11’* for A-K12. Interestingly, similar patterns were displayed by all Shigella phylons, featuring low richness and moderate to high entropy. Notably, a common theme across these phylons was the prevalence of *insG* as the dominant TE (**Fig 5b**).

The remaining 21% of genes in the rare genome result from TE insertions that fracture coding regions into smaller coding regions (Sheng et al. 2023), or horizontal gene transfer (HGT). As an example of HGT, a gene family found in the rare genome is *lapA*, encoding for a large adhesion pilus of over 6,000 amino acids in length, and commonly found in Pseudomonas species (see details in **SI**).

## Conclusions

Taxonomy going back to Linnaeus’ time was based on the form and phenotypic function of organisms. Then, classification based on genetics, such as certain alleles, emerged (e.g., multi-locus sequence tags (Jolley et al. 2018), and Clermont quadruplex (Clermont et al. 2013)). With whole-genome sequences becoming available, the MASH distance (Ondov et al. 2016) could be used to assess the relatedness of strains using genome-scale sequence similarity metrics. Now with genome sequences for a large number strains becoming available, we can annotate them and determine the species’ pangenome.

The pangenome is formally represented by the pangenome matrix, **P**, whose columns represent genomes and whose rows represent genes. Every column is then filled in with 0 or 1, the absence or presence of the gene in that genome, respectively. Since the number of sequenced genomes is now large, mathematical and machine learning methods can be applied to formulate global and rigorous classification schemes for a strain’s phylogeny based on the pangenome matrix. Such classification schemes are fundamental and will be at the root of bacterial taxonomy.

In this study we developed a classification schema based on the pangenome matrix using methods of machine learning. Remarkably, this approach gives a very clear definition of the gene content that differentiates strains that closely follows classical phylogroup definitions. The variably present genes populate the accessory genome, whose gene distribution amongst the strains can be used to obtain a mathematical definition of phylons, which are lists of genes that are found in the majority of the strains of a phylogroup.

This study enables a detailed, genomewide analysis of the genetic basis for the differential traits of the strains in the defined phylons and it gives a global multi-scale genetic structure of a species. This full exposé of the genetic composition of a bacterial species has many implications. With the availability of the alleleome (Catoiu et al. 2023), representing the global assessment of sequence variation of coding and intergenic regions, we can begin to understand the evolutionary history of a species and its phylons. The rare genome can keep track of horizontal gene transfer events and how they are assimilated into the species, and can provide new insights for understanding unique traits of a particular strain.

Detailing the phenotypic consequences based on the differential gene presence may take many years to fully resolve for all genetic traits of interest. This undertaking may have a fundamental effect on infectious disease. Reliable sequence-based rapid classification of pathogens isolated from a patient can accelerate physicians’ decision-making about pathogen identity and selection of treatment modalities.

One can anticipate that with a good coverage of genome sequences across the phylogenetic tree, we will be able to repeat the results of this study for larger and larger swaths of the tree. Perhaps, in the fullness of time, we will achieve a global genetic definition of the entire phylogenetic tree of bacteria.

## Methods

### Gathering and processing of sequence data from BV-BRC & NCBI RefSeq

We first downloaded the metadata of all genomes available on BV-BRC (Olson et al. 2023). Using this metadata file, we filtered out strains which were not *Escherichia coli* and those which only contained plasmid sequences. All “complete” sequences were further filtered by their L50 score (must equal 1) and their N50 score (greater than 4,000,000). Fragmented genomes were first filtered by their contig count, which was capped at 355 using previously defined metrics (Hyun et al. 2022). CheckM contamination (< 3.1%) and completeness (> 98.1%) scores were then used to filter fragmented genomes further (see code for more details on exact numerical thresholds chosen). The final collection was then downloaded from BV-BRC. This exact process was repeated for all *Shigella* strains and similarly for downloading *E. coli* strains from NCBI RefSeq. Genomes were then deduplicated and collated for further quality control (i.e. Mash filtration). In the end, only “complete” sequences were selected for pangenome analysis to ensure the pangenome had a gene presence/absence matrix (**P** matrix) of the highest quality.

### Genome annotation & pangenome generation

All downloaded genomes were re-annotated using prokka (Seemann 2014) for consistency in gene annotation when generating the pangenome. All re-annotated genomes were then screened by the *E. coli* PubMLST schema (Jolley et al. 2018) through the mlst github package (Seemann) to identify the sequence types for all strains in the pangenome. After this, the phylogroup for each strain was identified *in-silico* using the ClermonTyping github package (Beghain et al. 2018). Genomes were then collated to form a pangenome using CD-HIT (Li et al. 2001; Li et al. 2012). Gene families were identified using a sequence similarity and alignment cutoffs of 80% for both, as used in previous pangenome studies (Hyun et al. 2022). Once the pagenome was generated, all representative alleles which define a gene family as identified by CD-HIT were extracted and subjected to eggNOG gene annotation (Huerta-Cepas et al. 2019; Buchfink et al. 2021; Cantalapiedra et al. 2021). Genomes were also annotated for AMR gene annotation using the Resistance Gene Identifier tool (Alcock et al. 2023).

### Mash filtration & analysis

All downloaded genomes were run to generate pairwise Mash distance values. They were then separated into 6 groups: *Escherichia coli, Shigella sonnei, Shigella boydii, Shigella dysenteriae, Shigella flexneri*, and other *Shigella* species. For each group, the 99th percentile was calculated relative to the reference strain for each group and used as the filtration limit (that is, the top 1% of genomes in terms of Mash distance were filtered out for each group). Then, the Mash distance values were converted into Pearson correlation coefficients, which in turn were converted into Pearson correlation distances for Mash clustering, as outlined in Abram *et al* (Abram et al. 2021). A sensitivity analysis was performed to find the best threshold for clustering these values, which led to a value 0.1. Specifically, cutoff threshold for hierarchical clustering using seaborn’s inbuilt clustermap function was set to to various values until all major phylogroups of E coli were represented. This value (0.1) agreed with known domain knowledge, as Phylogroup C strains did not form their own cluster for any threshold value above 0.13 (Abram et al. 2021). This led to a total of 31 clusters, which was used to inform the rank of NMF decomposition.

### Defining the core, accessory, and rare genomes

The core, accessory, and rare genomes are defined using the cumulative gene distribution plot **(Fig 2b**) using methods outlined earlier (Hyun et al. 2022). Briefly, this gene plot forms an S-shaped curve and thus will always have an inflection point. The core genome is defined by taking the highest endpoint and traveling 90% of the distance from the inflection point to the endpoint. This corresponds with the elbow in the plot defining the core genes. A similar approach is used for defining the rare genome, except with the lowest endpoint instead of the highest one.

For the TEs analysis, transposable elements initially identified by PROKKA annotation were filtered. The start and end locations of each transposable element were determined on genomes across the entire pangenome. Genes that were not classified as transposable elements but had start and end locations within a transposable element were designated as passenger genes.

### TEs Richness and Evenness Calculation

A systematic genome sampling approach was implemented to ensure a representative and manageable dataset for this analysis. Phylons with fewer than 30 genomes were excluded from the analysis. For phylons having 30 or more genomes, we conducted a random sampling procedure to select 30 unique genomes.

This genome sampling strategy guaranteed that the dataset used for richness and evenness calculations was not only representative but also possessed a more even distribution. Consequently, it allowed for meaningful insights into TE diversity across various phylogroups. **Figure S5c** illustrates the sensitivity of TE richness, evenness, and count metrics to the random sampling of genomes.

Richness, representing the number of unique TEs within each phylon, was computed. Evenness, quantifying the distribution uniformity of TEs within a phylon, was calculated using the Shannon entropy formula.

### Non-negative Matrix Factorization

The scikit-learn implementation of NMF (Pedregosa et al. 2011) was used to perform the decomposition. NMF was run 50 times with a rank of 31 (derived from Mash clustering), an initialization of “nndsvd” (which generates sparser output matrices), and a maximum iteration limit of 5000 (the solution always converged before this limit was reached for all runs). The best run (as defined by the Frobenius norm, sum of squared residuals, and root-mean-square-error metrics) was selected for normalization. For each column in **L**, the 99th percentile was calculated and every value in that column was divided by this value, which ensured all but a few values were between 0 and 1. To ensure reconstruction consistency, the corresponding rows in the **A** matrix were multiplied by the same normalization values (see SI for more information). The **L** and **A** matrices were then binarized using k-means clustering (k=3), also implemented using scikit-learn. Each column of **L** was segregated into 3 clusters, and the genes in the cluster with the highest average mean were binarized to 1, with the genes in the other 2 clusters being set to 0. The same procedure was followed with binarizing the **A** matrix. This protocol ensured the threshold for binarization was always a conservative estimate. In some cases, this estimate in the **A** matrix was clearly too conservative (as evidenced by visualizing the histogram of strain affinities for each phylon), and in those cases, the threshold for binarization was manually lowered.

### Phylon Characterization

Given the large number of genes in the L matrix, phylons were initially characterized using the (binarized) **A** matrix. Phylons were first named based on the phylogroup of the strains with the highest affinity for each phylon, followed by the MLST value of these strains. If at least 90% of strains with high affinity were part of a particular Clermont-defined phylogroup, that phylon was mapped to that Clermont-defined phylogroup (e.g. if 90% of all strains in phylon2 map to phylogroup B2, that phylon is mapped to B2). In certain cases, the names of the phylons were changed to reflect well-known strains within the phylon (e.g. A-K12, A-BL21, etc.). Some phylons did not follow these patterns and were thus dubbed “uncharacterized” phylons. Strains with “high affinity” are defined as those strains which had an entry of 1 in the binarized **A** matrix for a particular phylon. For most strains, this only occurred once. For some strains, this occurred multiple times; in all cases, the strain had a high affinity for only one named phylon as the other high affinities were all for the uncharacterized phylons which consisted of mobile genetic elements.

## Code and data availability statement

All data (and code) pertaining to this study has been deposited onto Zenodo and can be found with this DOI: 10.5281/zenodo.10575748.

## Author Information

Siddharth M. Chauhan (SMC)

Omid Ardalani (OA)

Jason C. Hyun (JCH)

Jonathan M. Monk (JMM)

Patrick V. Phaneuf (PVP)

Bernhard O. Palsson (BOP)

## Author Contributions

Conceptualization: BOP & SMC, Data curation: SMC & PVP, Investigation: SMC & OA, Methodology: SMC, JMM, & JCH, Mentorship: JMM & BOP, Writing & Editing: All authors

## Acknowledgements

The authors would like to acknowledge Dr. Akanksha Rajput for useful discussions.This work was funded by the Novo Nordisk Foundation Grant Number NNF20CC0035580.

## Notes

https://zenodo.org/records/10575748

### Competing Interest Statement

The authors have declared no competing interest.

## References

Abram K, Udaondo Z, Bleker C, Wanchai V, Wassenaar TM, Robeson MS 2nd, Ussery DW. 2021. Mash-based analyses of Escherichia coli genomes reveal 14 distinct phylogroups. Commun Biol. 4(1):117. doi:10.1038/s42003-020-01626-5. http://dx.doi.org/10.1038/s42003-020-01626-5.

Achtman M, Zhou Z, Charlesworth J, Baxter L. 2022. EnteroBase: hierarchical clustering of 100 000s of bacterial genomes into species/subspecies and populations. Philos Trans R Soc Lond B Biol Sci. 377(1861):20210240. doi:10.1098/rstb.2021.0240. http://dx.doi.org/10.1098/rstb.2021.0240.

Alcock BP, Huynh W, Chalil R, Smith KW, Raphenya AR, Wlodarski MA, Edalatmand A, Petkau A, Syed SA, Tsang KK, et al. 2023. CARD 2023: expanded curation, support for machine learning, and resistome prediction at the Comprehensive Antibiotic Resistance Database. Nucleic Acids Res. 51(D1):D690–D699. doi:10.1093/nar/gkac920. http://dx.doi.org/10.1093/nar/gkac920.

Beghain J, Bridier-Nahmias A, Le Nagard H, Denamur E, Clermont O. 2018. ClermonTyping: an easy-to-use and accurate in silico method for Escherichia genus strain phylotyping. Microb Genom. 4(7). doi:10.1099/mgen.0.000192. http://dx.doi.org/10.1099/mgen.0.000192.

Biggel M, Xavier BB, Johnson JR, Nielsen KL, Frimodt-Møller N, Matheeussen V, Goossens H, Moons P, Van Puyvelde S. 2020. Horizontally acquired papGII-containing pathogenicity islands underlie the emergence of invasive uropathogenic Escherichia coli lineages. Nat Commun. 11(1):5968. doi:10.1038/s41467-020-19714-9. http://dx.doi.org/10.1038/s41467-020-19714-9.

Blattner FR, Plunkett G 3rd, Bloch CA, Perna NT, Burland V, Riley M, Collado-Vides J, Glasner JD, Rode CK, Mayhew GF, et al. 1997. The complete genome sequence of Escherichia coli K-12. Science. 277(5331):1453–1462. doi:10.1126/science.277.5331.1453. http://dx.doi.org/10.1126/science.277.5331.1453.

Borowiak M, Fischer J, Hammerl JA, Hendriksen RS, Szabo I, Malorny B. 2017. Identification of a novel transposon-associated phosphoethanolamine transferase gene, mcr-5, conferring colistin resistance in d-tartrate fermenting Salmonella enterica subsp. enterica serovar Paratyphi B. J Antimicrob Chemother. 72(12):3317–3324. doi:10.1093/jac/dkx327. http://dx.doi.org/10.1093/jac/dkx327.

Buchfink B, Reuter K, Drost H-G. 2021. Sensitive protein alignments at tree-of-life scale using DIAMOND. Nat Methods. 18(4):366–368. doi:10.1038/s41592-021-01101-x. http://dx.doi.org/10.1038/s41592-021-01101-x.

Cantalapiedra CP, Hernández-Plaza A, Letunic I, Bork P, Huerta-Cepas J. 2021. eggNOG-mapper v2: Functional Annotation, Orthology Assignments, and Domain Prediction at the Metagenomic Scale. Mol Biol Evol. 38(12):5825–5829. doi:10.1093/molbev/msab293. http://dx.doi.org/10.1093/molbev/msab293.

Catoiu EA, Phaneuf P, Monk J, Palsson BO. 2023. Whole-genome sequences from wild-type and laboratory-evolved strains define the alleleome and establish its hallmarks. Proc Natl Acad Sci U S A. 120(15):e2218835120. doi:10.1073/pnas.2218835120. http://dx.doi.org/10.1073/pnas.2218835120.

Clermont O, Christenson JK, Denamur E, Gordon DM. 2013. The Clermont Escherichia coli phylo-typing method revisited: improvement of specificity and detection of new phylo-groups. Environ Microbiol Rep. 5(1):58–65. doi:10.1111/1758-2229.12019. http://dx.doi.org/10.1111/1758-2229.12019.

Clermont O, Dixit OVA, Vangchhia B, Condamine B, Dion S, Bridier-Nahmias A, Denamur E, Gordon D. 2019. Characterization and rapid identification of phylogroup G in Escherichia coli, a lineage with high virulence and antibiotic resistance potential. Environ Microbiol. 21(8):3107–3117. doi:10.1111/1462-2920.14713. http://dx.doi.org/10.1111/1462-2920.14713.

Clermont Olivier, Bonacorsi Stéphane, Bingen Edouard. 2000. Rapid and Simple Determination of the Escherichia coli Phylogenetic Group. Appl Environ Microbiol. 66(10):4555–4558. doi:10.1128/AEM.66.10.4555-4558.2000. https://doi.org/10.1128/AEM.66.10.4555-4558.2000.

Cooper KK, Mandrell RE, Louie JW, Korlach J, Clark TA, Parker CT, Huynh S, Chain PS, Ahmed S, Carter MQ. 2014. Comparative genomics of enterohemorrhagic Escherichia coli O145:H28 demonstrates a common evolutionary lineage with Escherichia coli O157:H7. BMC Genomics. 15:17. doi:10.1186/1471-2164-15-17. http://dx.doi.org/10.1186/1471-2164-15-17.

Cuénod A, Agnetti J, Seth-Smith HMB, Roloff T, Wälchli D, Shcherbakov D, Akbergenov R, Tschudin-Sutter S, Bassetti S, Siegemund M, et al. 2023. Bacterial genome-wide association study substantiates papGII of Escherichia coli as a major risk factor for urosepsis. Genome Med. 15(1):89. doi:10.1186/s13073-023-01243-x. http://dx.doi.org/10.1186/s13073-023-01243-x.

Cuzon G, Naas T, Nordmann P. 2011. Functional characterization of Tn4401, a Tn3-based transposon involved in blaKPC gene mobilization. Antimicrob Agents Chemother. 55(11):5370–5373. doi:10.1128/AAC.05202-11. http://dx.doi.org/10.1128/AAC.05202-11.

Devarajan K. 2008. Nonnegative matrix factorization: an analytical and interpretive tool in computational biology. PLoS Comput Biol. 4(7):e1000029. doi:10.1371/journal.pcbi.1000029. http://dx.doi.org/10.1371/journal.pcbi.1000029.

Di Martino ML, Fioravanti R, Barbabella G, Prosseda G, Colonna B, Casalino M. 2013. Molecular evolution of the nicotinic acid requirement within the Shigella/EIEC pathotype. Int J Med Microbiol. 303(8):651–661. doi:10.1016/j.ijmm.2013.09.007. http://dx.doi.org/10.1016/j.ijmm.2013.09.007.

Dogan B, Suzuki H, Herlekar D, Sartor RB, Campbell BJ, Roberts CL, Stewart K, Scherl EJ, Araz Y, Bitar PP, et al. 2014. Inflammation-associated adherent-invasive Escherichia coli are enriched in pathways for use of propanediol and iron and M-cell translocation. Inflamm Bowel Dis. 20(11):1919–1932. doi:10.1097/MIB.0000000000000183. http://dx.doi.org/10.1097/MIB.0000000000000183.

Fang X, Monk JM, Mih N, Du B, Sastry AV, Kavvas E, Seif Y, Smarr L, Palsson BO. 2018. Escherichia coli B2 strains prevalent in inflammatory bowel disease patients have distinct metabolic capabilities that enable colonization of intestinal mucosa. BMC Syst Biol. 12(1):66. doi:10.1186/s12918-018-0587-5. http://dx.doi.org/10.1186/s12918-018-0587-5.

Fleischmann RD, Adams MD, White O, Clayton RA, Kirkness EF, Kerlavage AR, Bult CJ, Tomb JF, Dougherty BA, Merrick JM. 1995. Whole-genome random sequencing and assembly of Haemophilus influenzae Rd. Science. 269(5223):496–512. doi:10.1126/science.7542800. http://dx.doi.org/10.1126/science.7542800.

Huerta-Cepas J, Szklarczyk D, Heller D, Hernández-Plaza A, Forslund SK, Cook H, Mende DR, Letunic I, Rattei T, Jensen LJ, et al. 2019. eggNOG 5.0: a hierarchical, functionally and phylogenetically annotated orthology resource based on 5090 organisms and 2502 viruses. Nucleic Acids Res. 47(D1):D309–D314. doi:10.1093/nar/gky1085. http://dx.doi.org/10.1093/nar/gky1085.

Hyun JC, Monk JM, Palsson BO. 2022. Comparative pangenomics: analysis of 12 microbial pathogen pangenomes reveals conserved global structures of genetic and functional diversity. BMC Genomics. 23(1):7. doi:10.1186/s12864-021-08223-8. http://dx.doi.org/10.1186/s12864-021-08223-8.

Jolley KA, Bray JE, Maiden MCJ. 2018. Open-access bacterial population genomics: BIGSdb software, the PubMLST.org website and their applications. Wellcome Open Res. 3:124. doi:10.12688/wellcomeopenres.14826.1. http://dx.doi.org/10.12688/wellcomeopenres.14826.1.

Konstantinidis KT, Tiedje JM. 2005. Genomic insights that advance the species definition for prokaryotes. Proc Natl Acad Sci U S A. 102(7):2567–2572. doi:10.1073/pnas.0409727102. http://dx.doi.org/10.1073/pnas.0409727102.

Korea C-G, Badouraly R, Prevost M-C, Ghigo J-M, Beloin C. 2010. Escherichia coli K-12 possesses multiple cryptic but functional chaperone-usher fimbriae with distinct surface specificities. Environ Microbiol. 12(7):1957–1977. doi:10.1111/j.1462-2920.2010.02202.x. http://dx.doi.org/10.1111/j.1462-2920.2010.02202.x.

Kris A. Wetterstrand MS. 2019 Mar 13. The Cost of Sequencing a Human Genome. Genome.gov. [accessed 2023 Apr 18]. https://www.genome.gov/about-genomics/fact-sheets/Sequencing-Human-Genome-cost.

Larsonneur F, Martin FA, Mallet A, Martinez-Gil M, Semetey V, Ghigo J-M, Beloin C. 2016. Functional analysis of Escherichia coli Yad fimbriae reveals their potential role in environmental persistence. Environ Microbiol. 18(12):5228–5248. doi:10.1111/1462-2920.13559. http://dx.doi.org/10.1111/1462-2920.13559.

Lee DD, Seung HS. 1999. Learning the parts of objects by non-negative matrix factorization. Nature. 401(6755):788–791. doi:10.1038/44565. http://dx.doi.org/10.1038/44565.

Li W, Fu L, Niu B, Wu S, Wooley J. 2012. Ultrafast clustering algorithms for metagenomic sequence analysis. Brief Bioinform. 13(6):656–668. doi:10.1093/bib/bbs035. http://dx.doi.org/10.1093/bib/bbs035.

Li W, Jaroszewski L, Godzik A. 2001. Clustering of highly homologous sequences to reduce the size of large protein databases. Bioinformatics. 17(3):282–283. doi:10.1093/bioinformatics/17.3.282. http://dx.doi.org/10.1093/bioinformatics/17.3.282.

Lu S, Jin D, Wu S, Yang J, Lan R, Bai X, Liu S, Meng Q, Yuan X, Zhou J, et al. 2016. Insights into the evolution of pathogenicity of Escherichia coli from genomic analysis of intestinal E. coli of Marmota himalayana in Qinghai-Tibet plateau of China. Emerg Microbes Infect. 5(12):e122. doi:10.1038/emi.2016.122. http://dx.doi.org/10.1038/emi.2016.122.

Monk JM, Charusanti P, Aziz RK, Lerman JA, Premyodhin N, Orth JD, Feist AM, Palsson BØ. 2013. Genome-scale metabolic reconstructions of multiple Escherichia coli strains highlight strain-specific adaptations to nutritional environments. Proc Natl Acad Sci U S A. 110(50):20338–20343. doi:10.1073/pnas.1307797110. http://dx.doi.org/10.1073/pnas.1307797110.

Monk JM, Lloyd CJ, Brunk E, Mih N, Sastry A, King Z, Takeuchi R, Nomura W, Zhang Z, Mori H, et al. 2017. iML1515, a knowledgebase that computes Escherichia coli traits. Nat Biotechnol. 35(10):904–908. doi:10.1038/nbt.3956. http://dx.doi.org/10.1038/nbt.3956.

Nordmann P, Dortet L, Poirel L. 2012. Carbapenem resistance in Enterobacteriaceae: here is the storm! Trends Mol Med. 18(5):263–272. doi:10.1016/j.molmed.2012.03.003. http://dx.doi.org/10.1016/j.molmed.2012.03.003.

Norsigian CJ, Kavvas E, Seif Y, Palsson BO, Monk JM. 2018. iCN718, an Updated and Improved Genome-Scale Metabolic Network Reconstruction of Acinetobacter baumannii AYE. Front Genet. 9:121. doi:10.3389/fgene.2018.00121. http://dx.doi.org/10.3389/fgene.2018.00121.

O’Leary NA, Wright MW, Brister JR, Ciufo S, Haddad D, McVeigh R, Rajput B, Robbertse B, Smith-White B, Ako-Adjei D, et al. 2016. Reference sequence (RefSeq) database at NCBI: current status, taxonomic expansion, and functional annotation. Nucleic Acids Res. 44(D1):D733–45. doi:10.1093/nar/gkv1189. http://dx.doi.org/10.1093/nar/gkv1189.

Olson RD, Assaf R, Brettin T, Conrad N, Cucinell C, Davis JJ, Dempsey DM, Dickerman A, Dietrich EM, Kenyon RW, et al. 2023. Introducing the Bacterial and Viral Bioinformatics Resource Center (BV-BRC): a resource combining PATRIC, IRD and ViPR. Nucleic Acids Res. 51(D1):D678–D689. doi:10.1093/nar/gkac1003. http://dx.doi.org/10.1093/nar/gkac1003.

Ondov BD, Treangen TJ, Melsted P, Mallonee AB, Bergman NH, Koren S, Phillippy AM. 2016. Mash: fast genome and metagenome distance estimation using MinHash. Genome Biol. 17(1):132. doi:10.1186/s13059-016-0997-x. http://dx.doi.org/10.1186/s13059-016-0997-x.

Pedregosa F, Varoquaux G, Gramfort A, Michel V, Thirion B, Grisel O, Blondel M, Prettenhofer P, Weiss R, Dubourg V, et al. 2011. Scikit-learn: Machine Learning in Python. J Mach Learn Res. 12(85):2825–2830. [accessed 2023 Nov 14]. http://jmlr.org/papers/v12/pedregosa11a.html.

Perna NT, Plunkett G 3rd, Burland V, Mau B, Glasner JD, Rose DJ, Mayhew GF, Evans PS, Gregor J, Kirkpatrick HA, et al. 2001. Genome sequence of enterohaemorrhagic Escherichia coli O157:H7. Nature. 409(6819):529–533. doi:10.1038/35054089. http://dx.doi.org/10.1038/35054089.

Rajput A, Seif Y, Choudhary KS, Dalldorf C, Poudel S, Monk JM, Palsson BO. 2021. Pangenome Analytics Reveal Two-Component Systems as Conserved Targets in ESKAPEE Pathogens. mSystems. 6(1). doi:10.1128/mSystems.00981-20. http://dx.doi.org/10.1128/mSystems.00981-20.

Schauer R, Kamerling JP. 2018. Exploration of the Sialic Acid World. Adv Carbohydr Chem Biochem. 75:1–213. doi:10.1016/bs.accb.2018.09.001. http://dx.doi.org/10.1016/bs.accb.2018.09.001.

Seemann T. 2014. Prokka: rapid prokaryotic genome annotation. Bioinformatics. 30(14):2068–2069. doi:10.1093/bioinformatics/btu153. http://dx.doi.org/10.1093/bioinformatics/btu153.

Seemann T. mlst: :id: Scan contig files against PubMLST typing schemes. Github. [accessed 2023 Nov 14]. https://github.com/tseemann/mlst.

Seif Y, Kavvas E, Lachance J-C, Yurkovich JT, Nuccio S-P, Fang X, Catoiu E, Raffatellu M, Palsson BO, Monk JM. 2018. Genome-scale metabolic reconstructions of multiple Salmonella strains reveal serovar-specific metabolic traits. Nat Commun. 9(1):3771. doi:10.1038/s41467-018-06112-5. http://dx.doi.org/10.1038/s41467-018-06112-5.

Sheng Y, Wang H, Ou Y, Wu Y, Ding W, Tao M, Lin S, Deng Z, Bai L, Kang Q. 2023. Insertion sequence transposition inactivates CRISPR-Cas immunity. Nat Commun. 14(1):4366. doi:10.1038/s41467-023-39964-7. http://dx.doi.org/10.1038/s41467-023-39964-7.

The HC, Thanh DP, Holt KE, Thomson NR, Baker S. 2016. The genomic signatures of Shigella evolution, adaptation and geographical spread. Nat Rev Microbiol. 14(4):235–250. doi:10.1038/nrmicro.2016.10. http://dx.doi.org/10.1038/nrmicro.2016.10.

Varki A, Gagneux P. 2012. Multifarious roles of sialic acids in immunity. Ann N Y Acad Sci. 1253(1):16–36. doi:10.1111/j.1749-6632.2012.06517.x. http://dx.doi.org/10.1111/j.1749-6632.2012.06517.x.

Viladomiu M, Metz ML, Lima SF, Jin W-B, Chou L, JRI Live Cell Bank, Guo C-J, Diehl GE, Simpson KW, Scherl EJ, et al. 2021. Adherent-invasive E. coli metabolism of propanediol in Crohn’s disease regulates phagocytes to drive intestinal inflammation. Cell Host Microbe. 29(4):607–619.e8. doi:10.1016/j.chom.2021.01.002. http://dx.doi.org/10.1016/j.chom.2021.01.002.

Wirth T, Falush D, Lan R, Colles F, Mensa P, Wieler LH, Karch H, Reeves PR, Maiden MCJ, Ochman H, et al. 2006. Sex and virulence in Escherichia coli: an evolutionary perspective. Mol Microbiol. 60(5):1136–1151. doi:10.1111/j.1365-2958.2006.05172.x. http://dx.doi.org/10.1111/j.1365-2958.2006.05172.x.

Zhang H, Zong Z, Lei S, Srinivas S, Sun J, Feng Y, Huang M, Feng Y. 2019. A Genomic, Evolutionary, and Mechanistic Study of MCR-5 Action Suggests Functional Unification across the MCR Family of Colistin Resistance. Adv Sci. 6(11):1900034. doi:10.1002/advs.201900034. http://dx.doi.org/10.1002/advs.201900034.

