## Supplementary Information for "The Pangenome of *Escherichia coli*"

#### Supplementary Figures

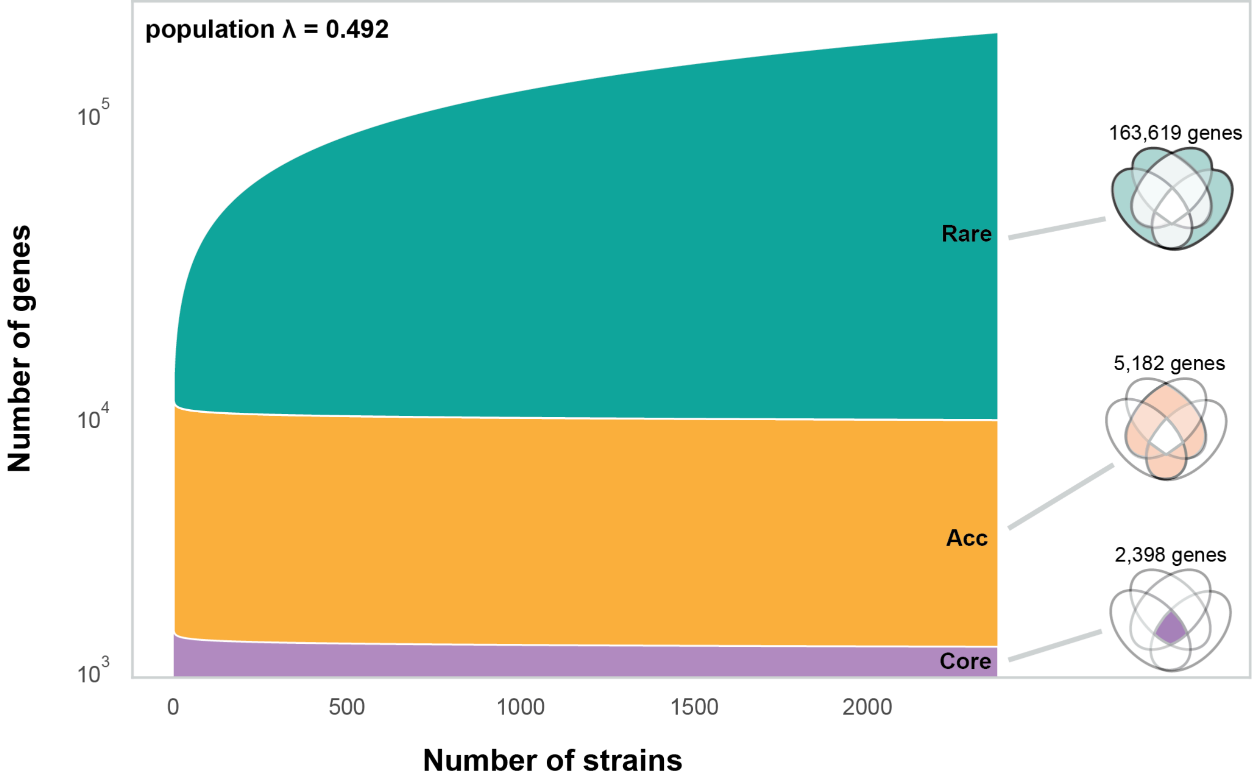


***Fig S1: A Heaps' Plot for the E coli pangenome.*** *The core genome (in purple) and the accessory genome (in orange) stabilize fairly quickly and do not change in gene content after a few hundred strains. Most of the growth in the* E. coli *pangenome comes from the gene number in the rare genome. It drives up the Heaps coefficient (λ) to a value to 0.492; Heap's Law is* $G=aN^{\lambda}$ *here G is the number of genes in the Pangenome (shown on the y-axis). N is the number of strains considered (shown on the x-axis), and 'a' and 'λ' are parameters obtained through curve fitting. When interpreting the rare genome size of 163,619 one needs to consider that 127,223 genes (79% of total) are variants of 315 transposon elements.*

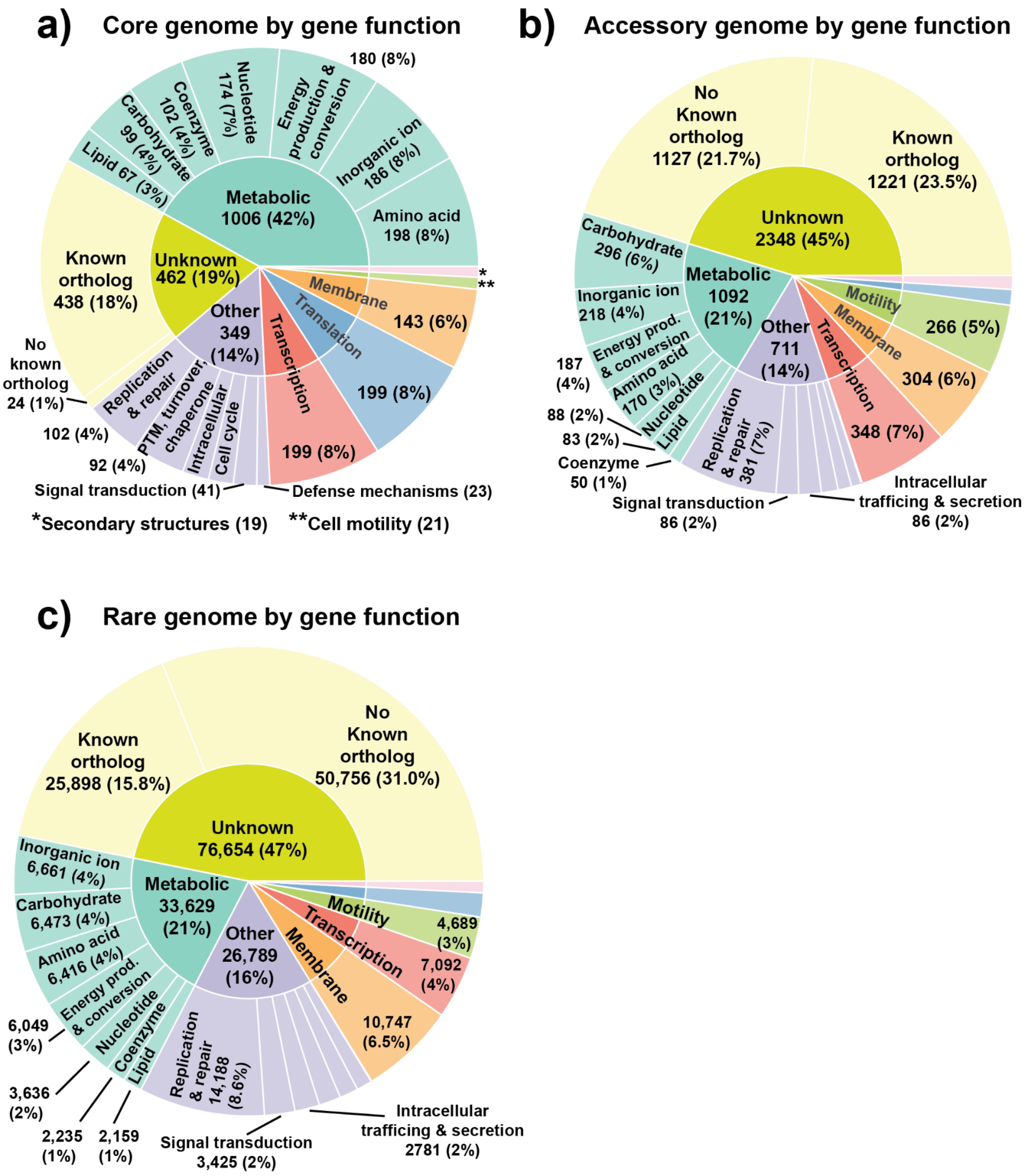


***Figure S2: Breakdown of core, accessory, and rare genomes by gene function:*** *The core genome is by far the most defined, with 81% of all its gene content having had their gene functions identified. The core genome is dominated by metabolic genes, which collectively form the core metabolism of Escherichia coli. The accessory and rare genomes, on the other hand, have roughly half of their gene portfolio with unknown gene functions. Transcription-, motility-, and membrane-related genes are also all present throughout the core, accessory, and rare genomes.* ***a)*** *A sunburst plot showcasing the breakdown of genes by function category in the core genome.* ***b)*** *A sunburst plot showcasing the breakdown of genes by function category in*

*the accessory genome.* ***c)*** *A sunburst plot showcasing the breakdown of genes by function category in the rare genome.*

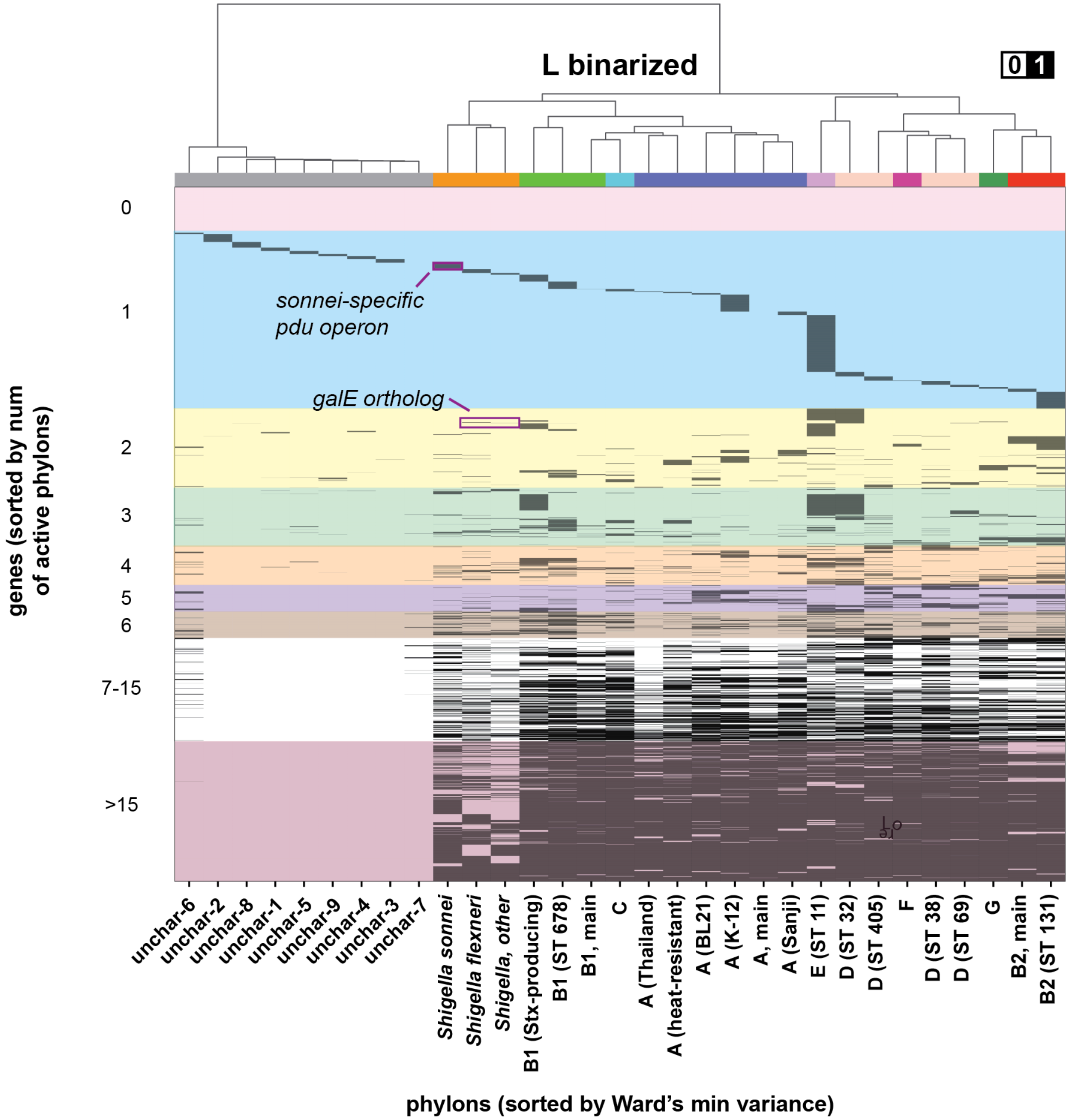


***Figure S3: The enlarged version of the binarized L matrix (from Fig 3d).*** *The rows represent genes and are ordered by the number of phylons a gene is found in. The location of two sets of genes that confer certain traits are highlighted (pdu is in one phylon, galE is found in two pylons). They are also highlighted in Figure 4.*

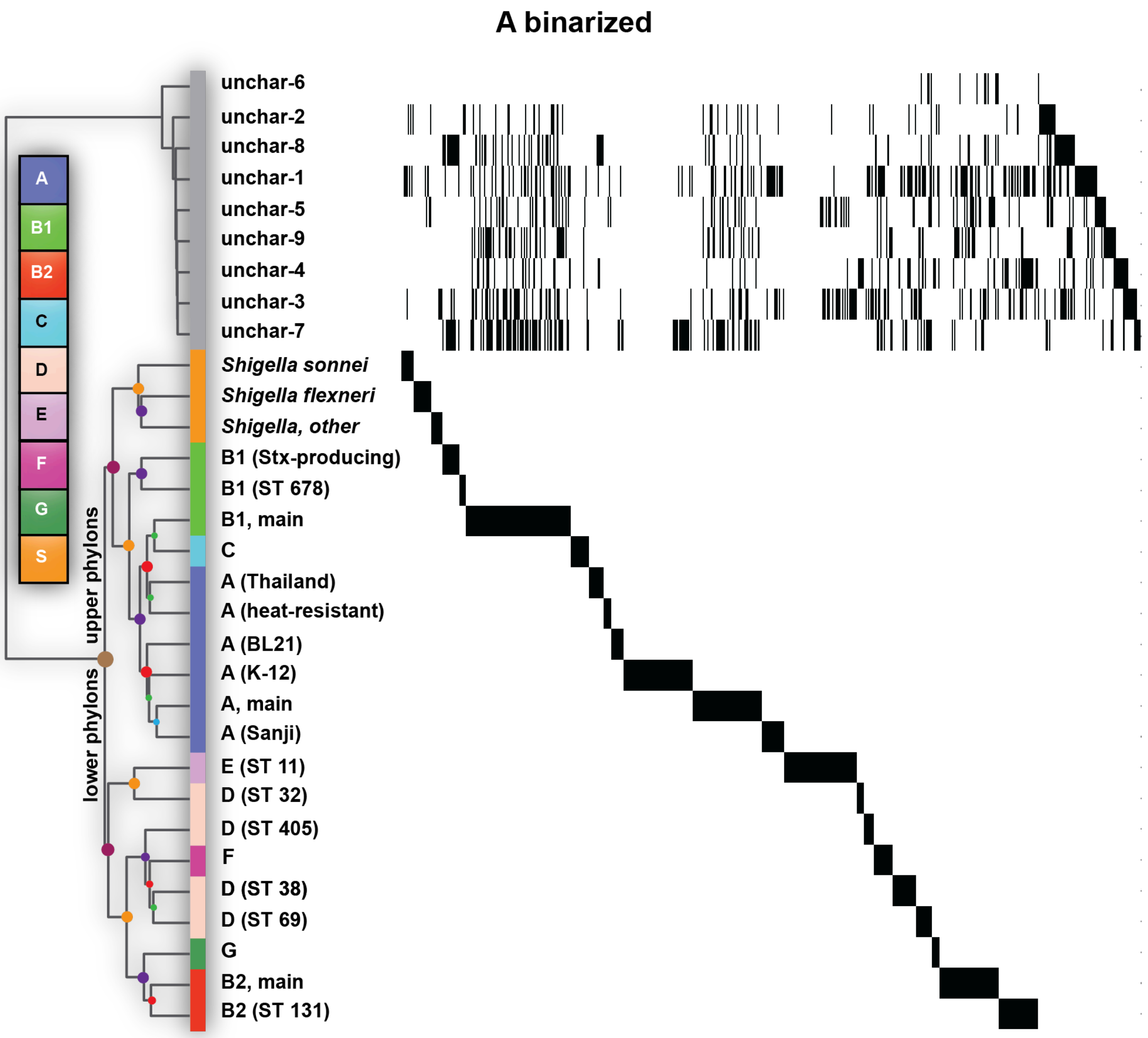


***Figure S4: The binarized A matrix demonstrating strain affinity for phylons.*** *Each row represents a phylon and each column represents a strain. The band structure seen across the 22 main phylons shows that all strains have a very strong affinity for a single main phylon and all other affinities are rounded down to zero when the matrix is binarized The upper part of the* ***A*** *matrix has a more complex structure as each of the 9 uncharacterized phylons have mobile elements that can be found in strains of all the phylogroups. The uncharacterized phylons consist of genes belonging to F-plasmids, sex pili, and other mobile genetic elements.*

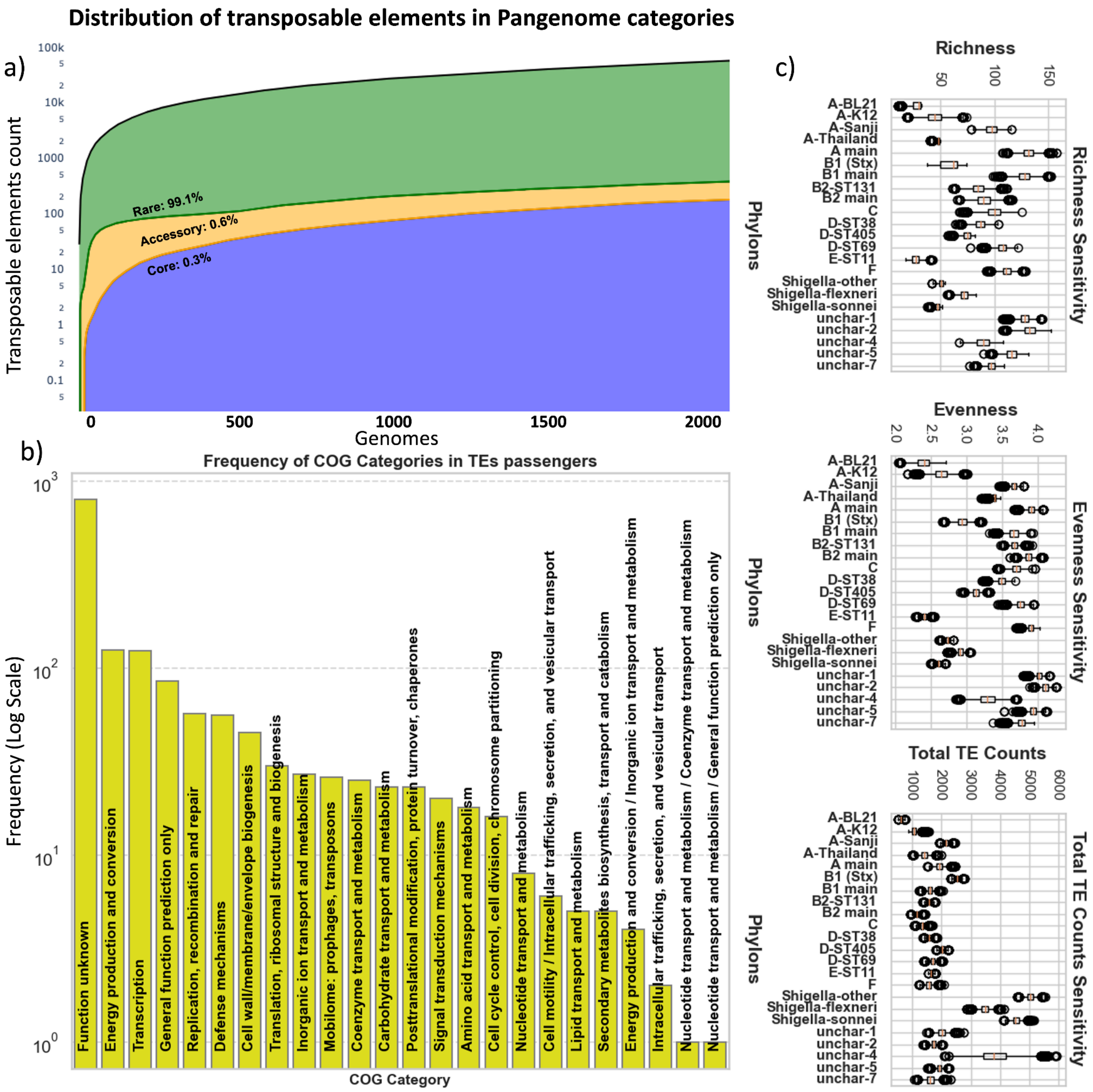


***Figure S5: Quantitative Analysis of Transposable Elements in Genomic Data a)*** *Illustrates the Heap's Law relationship in the dataset, showing the number of distinct transposable elements (TEs) (y-axis, log scale) against the number of genomes (x-axis). Each line represents a different TE category—core TEs (blue), accessory TEs (orange), and rare TEs (green)—with the area under each line shaded correspondingly.* ***b)*** *Bar plot visualizes the frequency of various COG categories in TEs passengers. Each bar represents a COG category, labeled directly on the bar, and the height of the bar indicates the frequency of that category on a logarithmic scale.* ***c)*** *Represent the results of sensitivity analysis, demonstrating the variability in transposable element richness, evenness, and total counts across different phylogroups. The analysis was conducted over 10,000 iterations, with each iteration involving a random selection of 30 genomes per phylogroup and the calculation of the corresponding TE metrics. The left panel shows the distribution of TE richness values, the middle panel depicts the evenness, and*

*the right panel illustrates the total TE counts for each phylogroup. The boxplots reveal the range and median of these metrics, highlighting their sensitivity to genome sampling.*

### Supplementary Text

**Defining the boundaries of the core, accessory, and rare genomes:** As mentioned in the main text, the exact boundaries between the core/accessory and the accessory/rare genomes do not affect the major conclusions of this study. For the core/accessory boundary, the addition of the core genomes into NMF does not fundamentally change the outputs we see, but rather creates a new "core-phylon" which consists of genes which are enriched across all strains. In fact, with the current boundaries, we already see 20 genes which are present in all phylons, with all of these genes being "sub-core" i.e. genes which are just under the threshold to be considered a core gene. Similarly, rare genes are so sparsely present that NMF cannot pick up their signal during decomposition, especially at a rank of 31, which is much lower than the column space of the P matrix. We also see this in Fig 3d, whereby 437 genes are not found in any of the phylons, with most of these genes being found between 7%-18% of all strains (right on or next to the accessory/rare genome boundary). Therefore, adding in more core genes or rare genes does not fundamentally affect the L matrix that is output from NMF decomposition.

**Transcriptome regulation of genes in the core genome:** There is information available on the expression patterns of the genes in the core genome. The transcriptome of the MG1655 strain has been modularized into sets of genes that are independently modulated, termed iModulons [53,54]. Of the 2255 core genes, 1259 (56%) are found in an iModulon.  Metabolism in *E. coli* is well characterized, and of the 1009 metabolic genes, 625 (62%) are found in an iModulon. Thus, the regulation of genes in the core genome that exhibit variable expression under more than 1000 growth conditions is documented [53,54], while many of the remaining genes seem to be constitutive.  A recent study found a similar number of proteins that seem to be constitutively depressed [55].

**Regarding the lapA gene found in the rare genome:** The specific strain this gene was found in (NCBI RefSeq: GCF_002854065.1) is called 14EC033 and is a clinical isolate collected from a patient in China in 2014. This strain, in addition to having a main chromosome, also contains seven other genomic elements labeled p14EC033a through p14EC033g, with each non-chromosome genomic element averaging 90 Kbp in length.

**Transposable Element analysis:** The TE sequences in our dataset were initially identified through prokka annotation. Subsequently, these TE sequences were categorized into 315 distinct groups based on their annotations. Additionally, CD-HIT clustering grouped these sequences into 2,920 separate clusters. The discrepancy in clustering results between CD-HIT and prokka can be attributed to the inherent diversity in TE structures. TEs are intricate constructs composed of various genes, random repeats, additional flanking elements, and, in some instances, fragments of broken genes. Furthermore, certain TEs exhibit additional components known as "passengers," further contributing to their diversity (notably, the top 40 groups among the 315 contained 773 unique passengers). CD-HIT clusters TEs based on sequence similarity, whereas prokka leverages specialized databases, such as ISfinder, tailored specifically for TEs to ascertain their identities.
